# Selective Plane Illumination Microscopy and Computing Reveal Differential Obliteration of Retinal Vascular Plexuses

**DOI:** 10.1101/2020.05.06.081463

**Authors:** Chih-Chiang Chang, Alison Chu, Scott Meyer, Michel M. Sun, Parinaz Abiri, Kyung In Baek, Varun Gudapati, Xili Ding, Pierre Guihard, Yichen Ding, Kristina I. Bostrom, Song Li, Lynn K. Gordon, Jie J. Zheng, Tzung K. Hsiai

## Abstract

Murine models of visual impairment provide micro-vascular insights into the 3-D network disarray in retinopathy. Current imaging and analysis tend to be confined to the 2-D retinal vasculature. We hereby integrated selective plane illumination imaging or known as light-sheet fluorescence microscopy (LSFM) with dual-illumination, followed by computational analyses, to reveal the topological network of vertical sprouts bridging the primary and secondary plexuses in a postnatal mouse model of oxygen-induced retinopathy (OIR). We revealed a preferential obliteration of the secondary plexus and bridging vessels despite a relatively unscathed primary plexus. We compared the local versus global vascular connectivity using clustering coefficients and Euler numbers, respectively. The global vascular connectivity in hyperoxia-exposed retinas was significantly reduced (*p* < 0.05, n = 5 vs. normoxia), whereas the local connectivity was preserved (*p* > 0.05, n = 5 vs. normoxia). We further applied principal component analysis (PCA) to automatically segment the vertical sprouts, corroborating the preferential obliteration of the interconnection between vertical sprouts and secondary plexuses that were accompanied with impaired vascular branching and connectivity, and reduced vessel volumes and lengths (*p* < 0.05, n=5 vs. normoxia). Thus, integration of 3-D selective plane illumination with computational analyses allows for early detection of global and spatially-specific vaso-obliteration, but preserved local reticular structure in response to hyperoxia-induced retinopathy.

## Introduction

Aberrant retinal angiogenesis is a hallmark of numerous retinal disorder-mediated vasculopathies, including retinopathy of prematurity and diabetic retinopathy, resulting in visual impairment^1,2^. Premature newborns are susceptible to hyperoxia and the development of retinopathy of prematurity as a result of supplemental oxygen therapy often needed due to premature development of their lungs, leading to vascular bed depletion^3-5^. These newborns further develop ischemia-induced neovascularization in the retina due to local hypoxia and increased metabolic demand^4,6^. Similarly, diabetic patients are also prone to developing proliferative retinopathy and aberrant neovascularization as a result of hypoxia^7,8^.

To quantify microvascular damage that occurs early in diabetic retinopathy, ophthalmologists have relied on several clinical imaging modalities, including fundus photography, fluorescein angiography (FA), and optical coherence tomography angiography (OCTA). These imaging modalities allow for screening and monitoring of vision-threatening complications in adults ^9-11^. OCT-angiography enables visualization of preclinical retinal vascular changes, including the remodeling of the foveal avascular zone (FAZ), capillary nonperfusion, and reduction of capillary density^12-16^. However, the quality of these clinical images in neonates and young children is often limited by ocular movement and restricted field of view.

To elucidate vasculopathy in the presence of retinopathy of prematurity, researchers have used the mouse models of oxygen-induced retinopathy (OIR) ^17,18^. Akin to ocular development in humans, the post-natal mouse retina is considered to be a viable model to underpin the development of retinal vascular network^1,2,17^. The existing modalities to image and quantify the retinal vascular network are frequently confined to either 2-D or whole-mount samples^19,20^. To identify the early abnormalities and progression of OIR in 3-D, we hereby integrated selective plane illumination microscopy or known as light-sheet fluorescence microscopy (LSFM) with dual-illumination, followed by quantitative analyses to acquire 3-D volumetric images^21-25^. Unlike confocal or two-photon microscopy which usually utilizes a point scanning approach, LSFM generates a sheet of light to rapidly scan across the optically-cleared specimens^21,26,27^. This approach provides an entry point to investigate the intact murine ocular system and rapid data acquisition to minimize photo-bleaching. LSFM further provides the high axial resolution needed to precisely quantify the multi-layered vascular network; namely, the vertical sprouts embedded between the primary and secondary plexuses ^1,17,28^.

To this end, we customized an optical clearing technique to preserve the 3-D conformation of the hemispherical murine retina and its vascular network for LSFM imaging. Following hyperoxia (75% O_2_) exposure to post-natal mice for 5 days (P7-P12), we quantified the preferential obliteration of vertical sprouts and secondary plexuses that peaked at day 12. Using principal component analysis (PCA), we developed an automated segmentation algorithm to demonstrate a significant reduction in the volume fraction of the vertical sprouts that interconnect between the primary and secondary plexuses. By using both Euler numbers and clustering coefficients, we revealed a reduction in the global vascular connectivity, but the preserved local connectivity in oxygen-induced retinopathy (OIR). With the copious data generated from LSFM techniques, our computational analysis, including PCA, was applied to identify deep capillary obliteration underlying OIR-mediated retinopathy outcomes. We hereby establish a novel multi-scale pipeline to unravel preferential obliteration of secondary plexus and vertical sprouts in the postnatal murine OIR model, allowing for detection of early changes in hyperoxia-mediated microvascular disarray underlying progression to visual impairment.

## Results

### Schematic pipeline for dual-illumination light-sheet imaging of the mouse retinal vasculature

To achieve high-spatial resolution for the 3-D complicated retinal vascular network (Fig 1A-B), we developed an eight-step pipeline for light-sheet imaging and quantification (Fig. 1C). Details will be introduced in the following sections. After euthanizing the mice at postnatal day 12 (P12), the ocular globe was enucleated and dissected to obtain the intact hemispherical retinas followed by polymerization and lipid removal in the monomer solution and clearing solution for optical transparency (Fig. 1D). Next, immunofluorescence staining with Isolectin B4 (Vector Lab, CA) was performed to label the vascular endothelium in preparation for light-sheet imaging. Following filament tracing, the morphology and topology of the retinal vasculature were characterized by quantifying the vascular branching points, Euler characteristics, and clustering coefficients.

**Figure 1.**
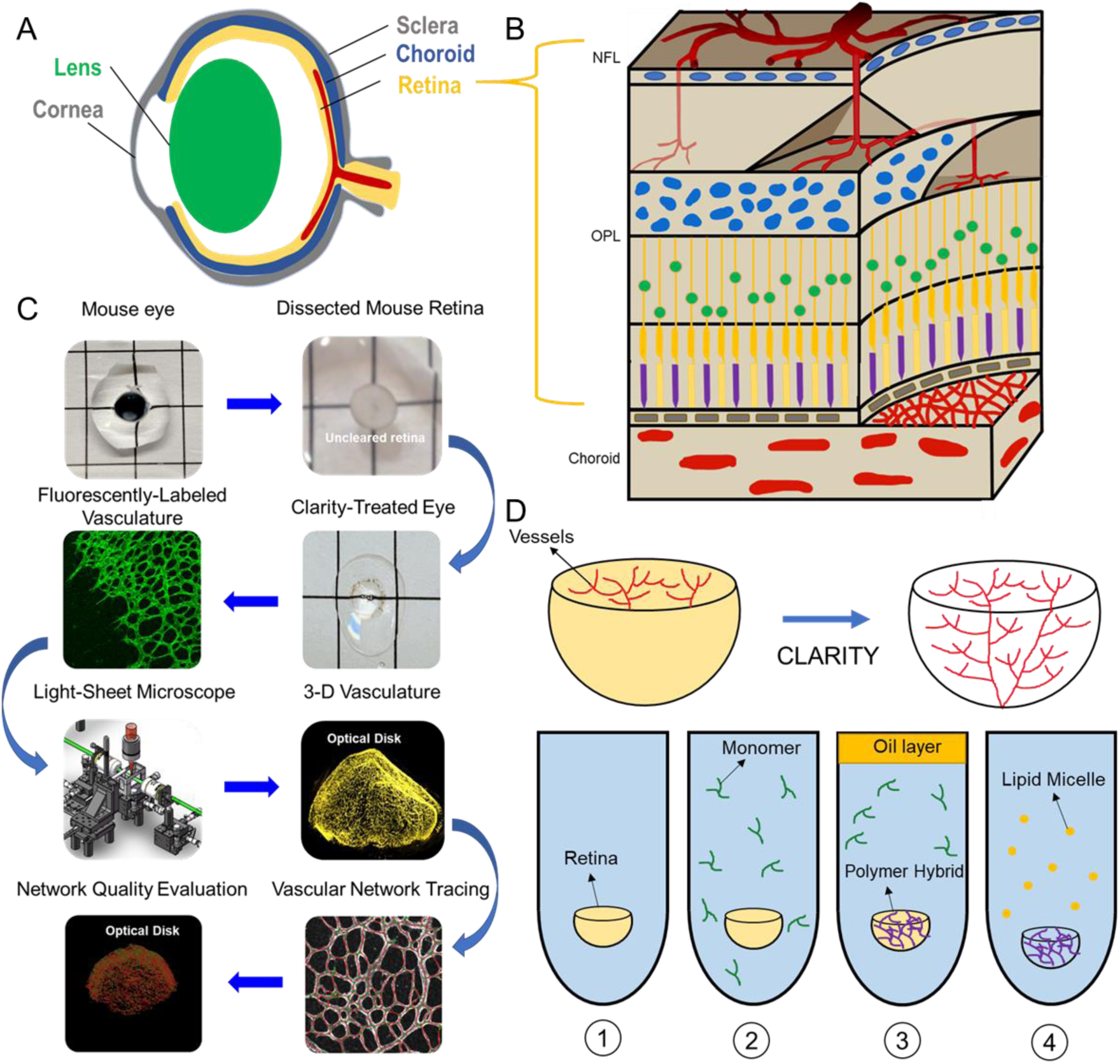
Light-sheet fluorescence microscopy (LSFM) to uncover the 3-D micro-vascular network. **(A)** A schematic illustration of the intact ocular globe, consisting of retinal and choroidal vasculature that were imaged and analyzed. **(B)** A superficial primary retinal vascular plexus lies in the nerve fiber layer (NFL), whereas the secondary plexus is located deep in the outer plexiform layer (OPL). **(C)** The pipeline quantitatively analyzed the 3-D hemispherical retinal vascular plexus using modified CLARITY and LSFM. **(D)** Modified passive CLARITY was applied to optically clear the retina as articulated in the Methods section.

### 3-D characterization of the vascular network under normoxia condition

LSFM with dual-illumination was employed to unravel the 3-D vascular network in the hemispherical retina in which the endothelial cells were labeled with Alexa Fluor 488 (Fig. 2A-B). Representative volumes of interest (VOIs) were selected, including the 2-D section (red dashed line), peripheral vascular networks in the peripheral region (orange box), the middle region (blue box) (Fig. 2B), and 2-D slice (Fig. 2C), to highlight the two-layered plexus and its 2-D connections. The LSFM-acquired 3-D images provide high-axial resolution as demonstrated by the color-coded depth penetration into the hemispherical retinal vasculature (Fig. 2C). The volumetric rendering of the VOI (blue box) revealed the 3-D vascular sprouts interconnected with the primary and secondary plexuses (Fig. 2D, Supplementary Movie 1). The light-sheet imaging further allowed for visualization of the 3-D vascular network from the selected VOIs at various depths (Fig. 2E-F), uncovering the interconnected capillary vascular bed at 5 μm in diameter via the color-coded depth imaging (Fig. 2G-H). Thus, our custom-built light-sheet microscope with dual illumination revealed the hemispherical vascular network with high axial resolution to unravel the novel multi-layered capillary plexuses under normoxia condition.

**Figure 2.**
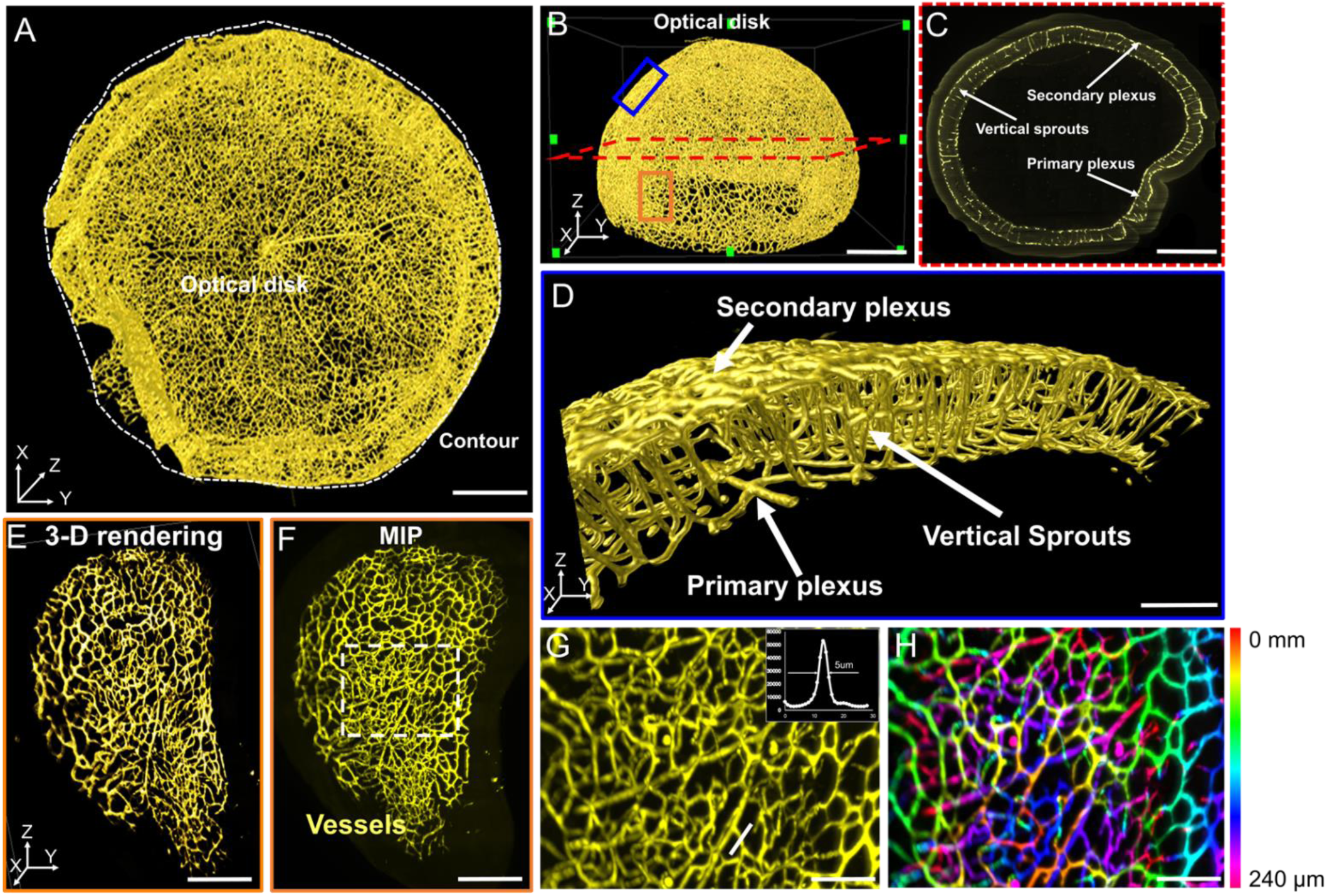
LSFM imaging of the intact 3-D hemispherical retina. **(A-B)** The intact vasculature in the 3-D retina provides representative regions of interest. The white dashed line depicts the contour of the hemispherical retina. **(C)** The vertical sprouts, bridging the primary and secondary plexus, are highlighted in the representative 2-D section (red dashed lines in **B**) of the retinal vascular network. **(D)** 3-D vertical sprouts between the primary and secondary vascular plexus (blue box in **B**) are located in the nerve fiber layer and the outer plexiform layer. **(E)** The 3-D and **(F)** 2-D peripheral regions (orange box in **B**) of the retina are accentuated. **(G)** The diameter of the capillary is ∼ 5-10 µm and **(H)** the 3-D capillary network (dashed box in **F**) is color-coded to indicate the depth. Scale bars: (A-C) 500 µm; (D) 200 µm; (E-F) 300 µm; (G-H) 50 µm. Depth color-coded scale: 0∼240 µm in (H).

### Morphological and topological quantification of the 3-D retinal vascular network under normoxia condition

We first applied the filament tracing strategy to characterize the retinal morphology and topology. Vascular connectivity and cluster coefficients were quantified (Fig. 3A, Supplementary Movie 2), as illustrated by the nodes (in green) interconnecting with the 3-D vascular network (in red). The connectivity and mean clustering coefficients for the intact retina in the absence of hyperoxia exposure were 1.6×10^4^ and 2.9×10^−2^, respectively. The former coefficient indicates the high number of loops present in the retinal vascular network, and the latter denotes the low degree of clustering of these individual nodes. We interrogated the central (yellow disk), middle (orange donuts), and peripheral (blue donuts) regions of the hemispherical network (Fig. 3). We selected 5 volumes of interest (VOIs) from each of the regions for analysis as a standard assessment for retinopathy ^11^. The 3-D vascular structure was visualized (Fig. 3B, D, and F), allowing for the application of filament tracing to characterize the different vessel lengths, branching points, and connections from the central, middle to peripheral regions. Using the filament tracing results, we were able to label the nodes in blue and segments in green (Fig. 3C, E, and G). The means of the entire vessel lengths were 6.62×10^3^ ±7.4×10^2^ µm for the central, 7.29 x10^3^ ±8.9×10^2^ µm for the middle, and 8.26 x10^2^ ±1.9×10^3^ µm for the peripheral regions. The difference among these 3 regions was statistically insignificant in the absence of hyperoxia exposure (*p* > 0.05, n=5). The means of the entire vessel volumes were 9.19 x 10^5^ ±1.63×10^5^ µm^3^, 9.81 x 10^5^ ±1.59×10^5^ µm^3^, and 9.27 x 10^5^ ±2.5×10^5^ µm^3^, respectively. There were 146 ±23.1 nodes and 32.3±9.3 in connectivity in the central region, 157.3 ±41.3 nodes and 35 ±9.85 in connectivity in the middle region, and 172.3 ±49.1 nodes and 43.3 ±16.1 in connectivity in the peripheral region (Fig. 3H-K). Thus, the differences in vascular lengths, volumes, connections, and branching points among the three representative regions of the retinal vasculature were statistically insignificant under the normoxia condition.

**Figure 3.**
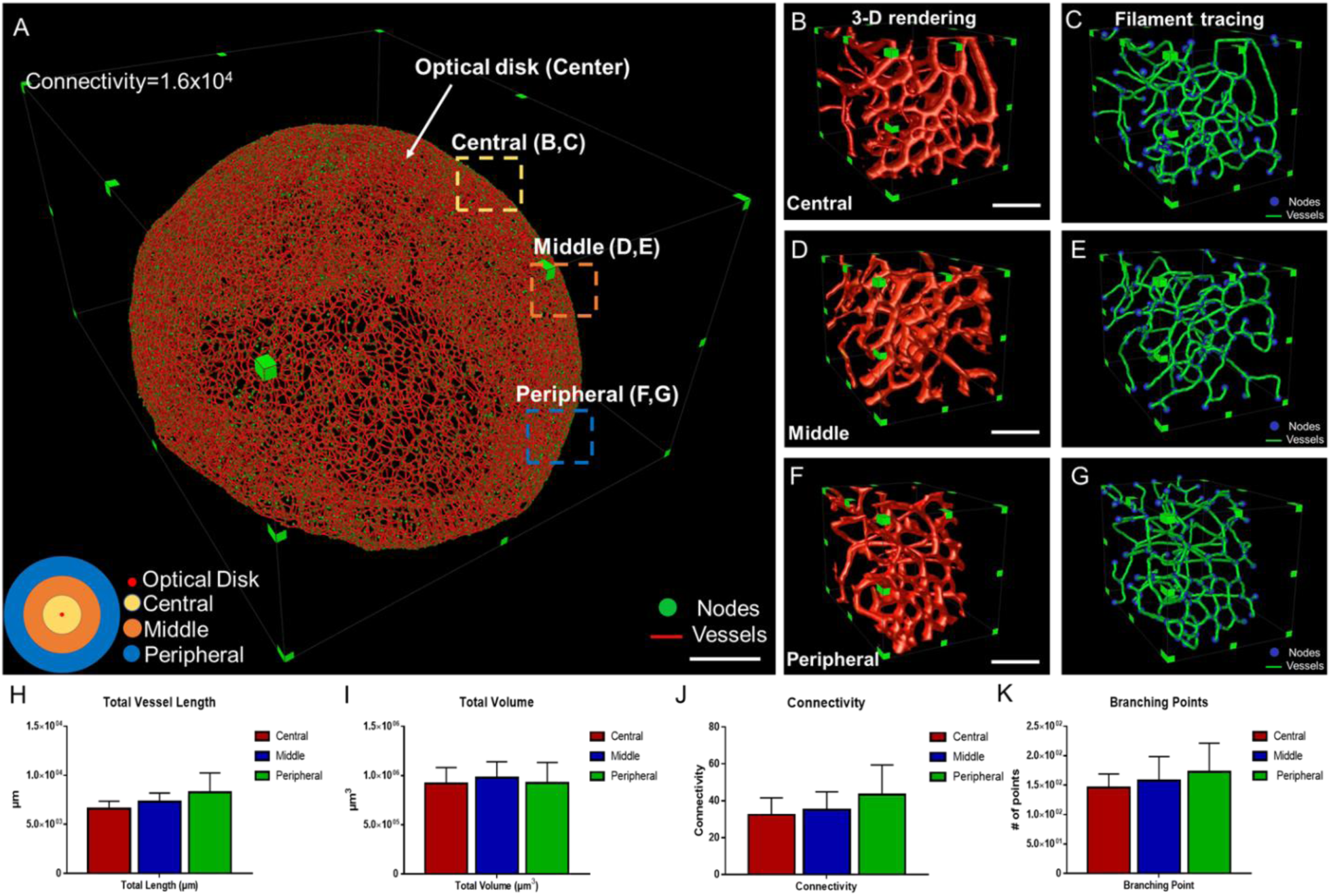
The quantitative comparison between the regions of healthy vascular network. **(A)** 3-D filament tracing of retinal vasculature was performed from the intact hemisphere retina. Representative **(B, D, F)** 3-D rendering and **(C, E, G)** filament tracing of different volumes of interest (VOI) were demonstrated in the central (B, C), middle (D, E) and peripheral (F, G) regions of the intact retina. **(H-K)** Quantification of the morphological and topological parameters; vessel lengths, vessel volumes, connectivities, and branching points for different VOIs from the retina, demonstrating the capability and flexibility of our pipeline to quantify the specific region of interest. Scale bar: (A) 500 µm; (B-G) 100 µm.

### Changes in morphological and topological quantification under hyperoxia condition

In response to hyperoxia, the 3-D vascular network (Fig. 4A-B, Supplementary Movie 3-4) and filament tracing (Fig. 4C, Supplementary Movie 5) in the oxygen-induced retinopathy (OIR) group were distinct from the normoxia groups (Fig. 4). A preferential depletion of the capillaries in the primary plexus developed in the central region, whereas depletion of vertical sprouts and secondary plexus occurred in all three regions as evidenced by the selected VOI (Fig. 4D-E). We further compared the topological parameters (Fig. 4F), including total length (normoxia: 1.2×10^6^±3.1×10^4^; OIR:2.6×10^5^±1.3×10^4^ µm, *p*<0.001, n=5), total volume (normoxia: 7.3×10^7^ ±1.4×10^7^; OIR: 3.2×10^7^±8.2×10^6^ µm^3^, *p*<0.05, n=5), branching point (normoxia: 2.7×10^4^±9.6×10^2^; OIR: 5.8×10^3^±1.3×10^3^, *p*<0.001, n=5) and connectivity (normoxia:1.1×10^4^±6.9×10^2^; OIR: 2.3×10^3^ ±9.1×10^2^, *p*<0.01, n=5) for the intact retinal vascular network (Fig. 4F). Taken together, our quantitative analyses corroborate the preferential obliteration of the 3-D capillary network in response to hyperoxia.

**Figure 4.**
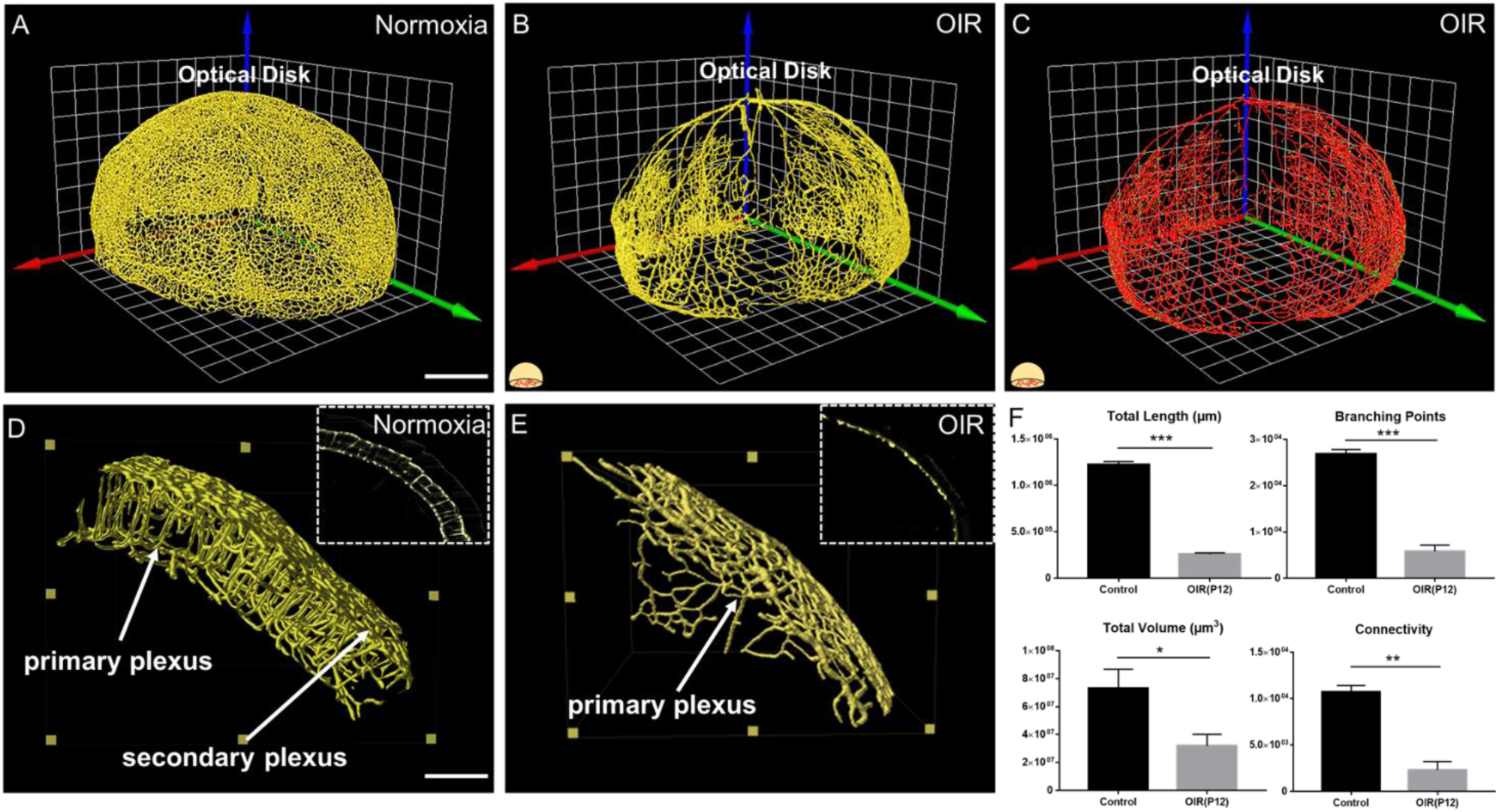
3-D vascular network to highlight the preferential vaso-obliteration in vertical sprouts and the secondary plexus in OIR. 3-D rendering of retinal vasculature was performed from the **(A)** normoxia and **(B)** OIR intact retina, demonstrating marked vaso-obliteration in the OIR retina. **(C)** 3-D filament tracing of vasculature for the intact retina was demonstrated from the OIR group. **(D-E)** 3-D rendering results from two volumes of interest (VOIs) in the normoxia (D) and OIR (E) groups indicating the differential depletion and absence of secondary plexus and vertical sprouts in the OIR mice. **(F)** Quantification of the morphological and topological parameters revealed the statistically significant lower values of total lengths, total volumes, connectivities and branching points in the OIR group. Scale bar: (A-C) 500 µm; (D-E) 100 µm.

### Frequency of clustering coefficients to demonstrate local vs. global vascular network

We computed the distribution of the clustering coefficients for each node in the vascular network (Fig. 5A-B). Clustering coefficients represent the degree of regional vascular clustering, ranging from 0 to 1. In both normoxia-(P12) and hyperoxia-treated mice, approximately 10% of the nodes exhibited positive clustering coefficients, whereas 90% of the nodes were nearly zero (grey), suggesting a paucity of connection with neighboring nodes in the 3-D vasculature. The common clustering coefficients and frequent degrees were 0 and 3, respectively, in the vascular network (Fig. 5B-C). These findings indicate that most of the nodes form connections with three neighboring nodes with no connections between its neighbors, supporting the reticulated structure of the 3-D vascular network, analogous to a honeycomb pattern. In addition, the local reticulated pattern was preserved despite hyperoxia-induced depletion of capillaries. We further compared the average clustering coefficients among the central, middle, and peripheral regions of the retina (Fig. 6A-B), revealing statistically insignificant differences between normoxia and hyperoxia conditions (central = 0.026±0.012 vs. 0.047±0.032, *p* > 0.05; middle = 0.027±0.018 vs. 0.039±0.013, *p* > 0.05; peripheral = 0.044±0.019 vs. 0.038±0.018, *p* > 0.05, n = 5). This comparison revealed that the local connectivity was preserved, whereas the global connectivity was significantly reduced (local = 0.032±0.018 vs. 0.043 ±0.024, *p* > 0.05, n = 5 and global = 1.1×10^4^±6.9×10^2^ vs. 2.3×10^3^±9.1×10^2^, *p* < 0.05, n = 5) (Fig. 4 and 6). Thus, clustering coefficients support the preserved local, but impaired global capillary connection in response to hyperoxia.

**Figure 5.**
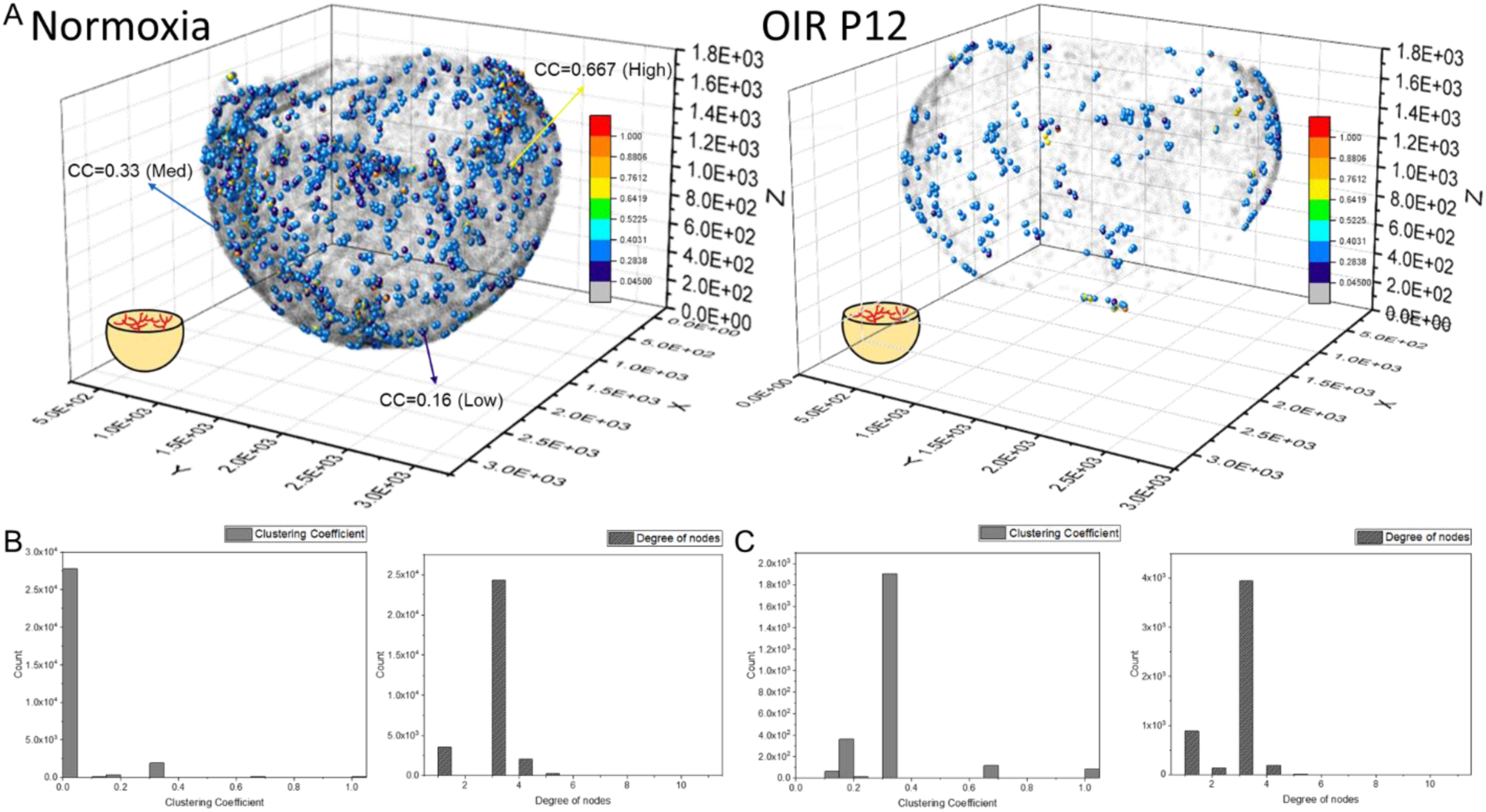
The quantification of the clustering coefficients for the intact vasculature. **(A)** 3-D scatter plot of the clustering coefficients reveals all branching nodes and terminal nodes traced. ∼90% of the nodes have a cluster coefficient of zero (Grey nodes). ∼10% of the nodes have clustering coefficients ranging from 0.05 to 1 in both normoxia and hyperoxia groups. **(B)** The histogram of clustering coefficients and degree of nodes demonstrates that the majority of the nodes have three connecting neighbors with a paucity of neighbor connection in the normoxia group. **(C)** Despite the significant reduction of the global connectivity and number of nodes, the OIR group demonstrated the same trend in local patterns as the normoxia group, with the majority of the nodes sharing three connecting neighbors with a paucity of the neighbor connection.

**Figure 6.**
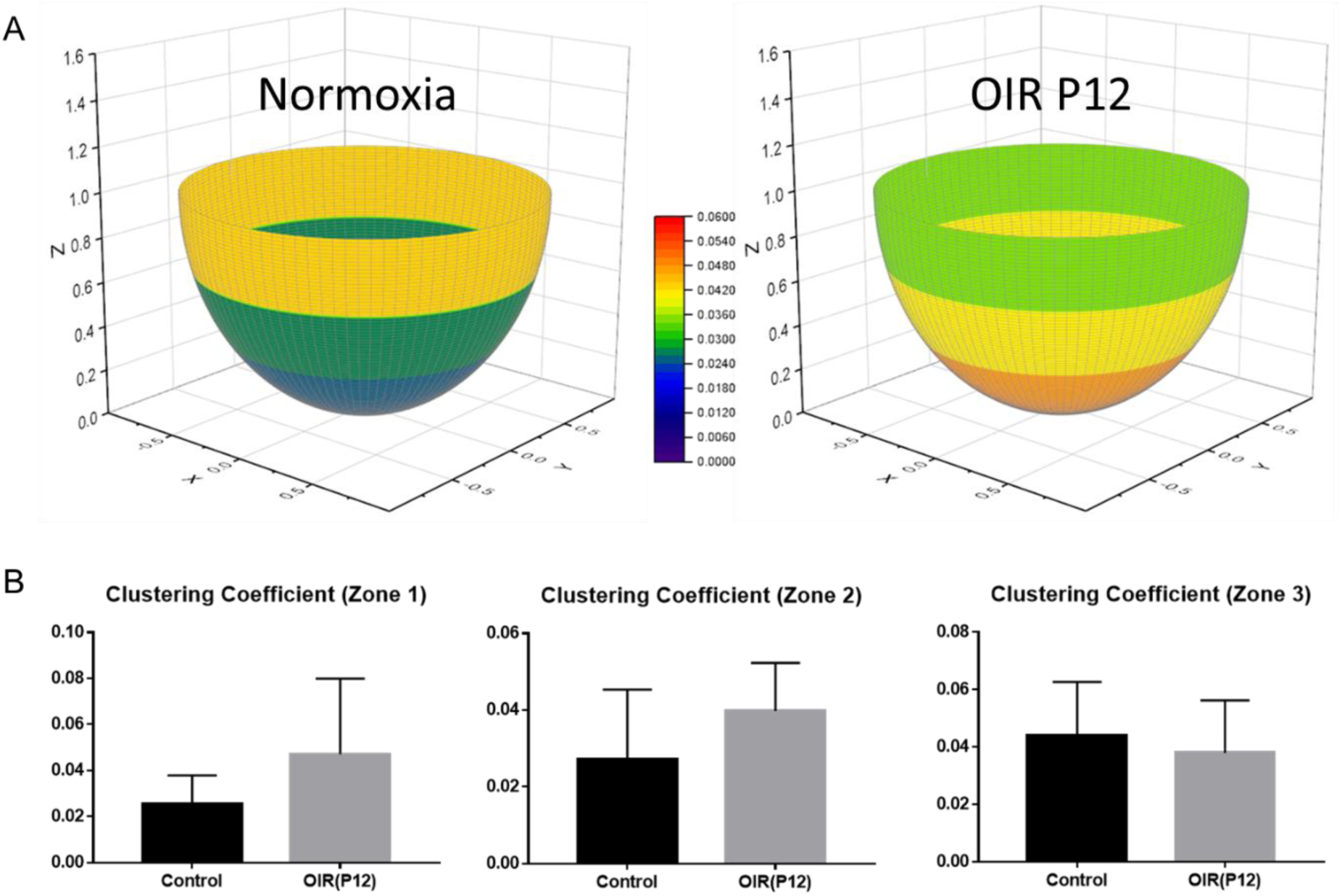
Average clustering coefficients in different regions of the retina. **(A)** The retinas were divided into three regions (central, middle, peripheral) as previously described. The average clustering coefficients of three individual regions were calculated and compared between the two groups. **(B)** The average clustering coefficients indicate no statistically significant difference among all three regions in both groups, supporting the notion that the local connection and basic structure of the network are preserved despite vaso-obliteration in OIR retinas (n=5 per group).

### Computational analyses of the plexuses and vertical sprouts following hyperoxia condition

Segmented 3-D image stacks for the entire retinal vasculature were derived from filament tracing results to construct an intact retinal vasculature structure. We randomly selected 4 VOIs with 200 µm in thickness to quantify the volume fraction of vertical sprouts, defined as the volume of vertical sprouts divided by the total volume of the vascular network. The representative images for the maximum intensity projection (MIP) of the automated segmented plexus and vertical sprouts were assessed from selected image slices (Supplementary Figure 1B-C). In addition, the 3-D rendering results (Fig. 7A-B) of the plexus (in yellow) and vertical sprouts (in blue) demonstrate the distinct phenotype and statistically significant obliteration (*p*<0.05 vs. normoxia, n=5) of the vertical sprouts following hyperoxia treatment (Fig. 7A & B). Differential obliteration was observed; namely, 17.98%±5.2% vs. 0.69%±0.75% in the middle, and 18.89%±7.43% vs. 0.51%±0.62% in the peripheral regions (Fig. 7C-D). To validate our quantification, we segmented the vertical sprouts and plexuses via manual and automated labeling. The generalized dice coefficients were calculated to compare the similarity between the manual annotation and the automated segmentation under different conditions^29^. To optimize automated segmentation, we introduced the sliding window (40×40×40 µm) with 20 µm in shifting distance and 35 degrees as the cutoff angles (Fig. 7E). Maximum dice coefficient and low standard deviation indicate accurate segmentation of vertical sprouts (Supplementary Tables 1-3). Finally, the representative VOI demonstrated the color-coded (X: Blue, Y: Green: Z: Red) 3-D rendering of the vascular networks based on the orientation of the vessels (Fig. 7F). Our computational strategy allows for multi-scale interrogation of the entire 3-D vascular network (macro-scale) vs. small VOIs (micro-scale).

**Figure 7.**
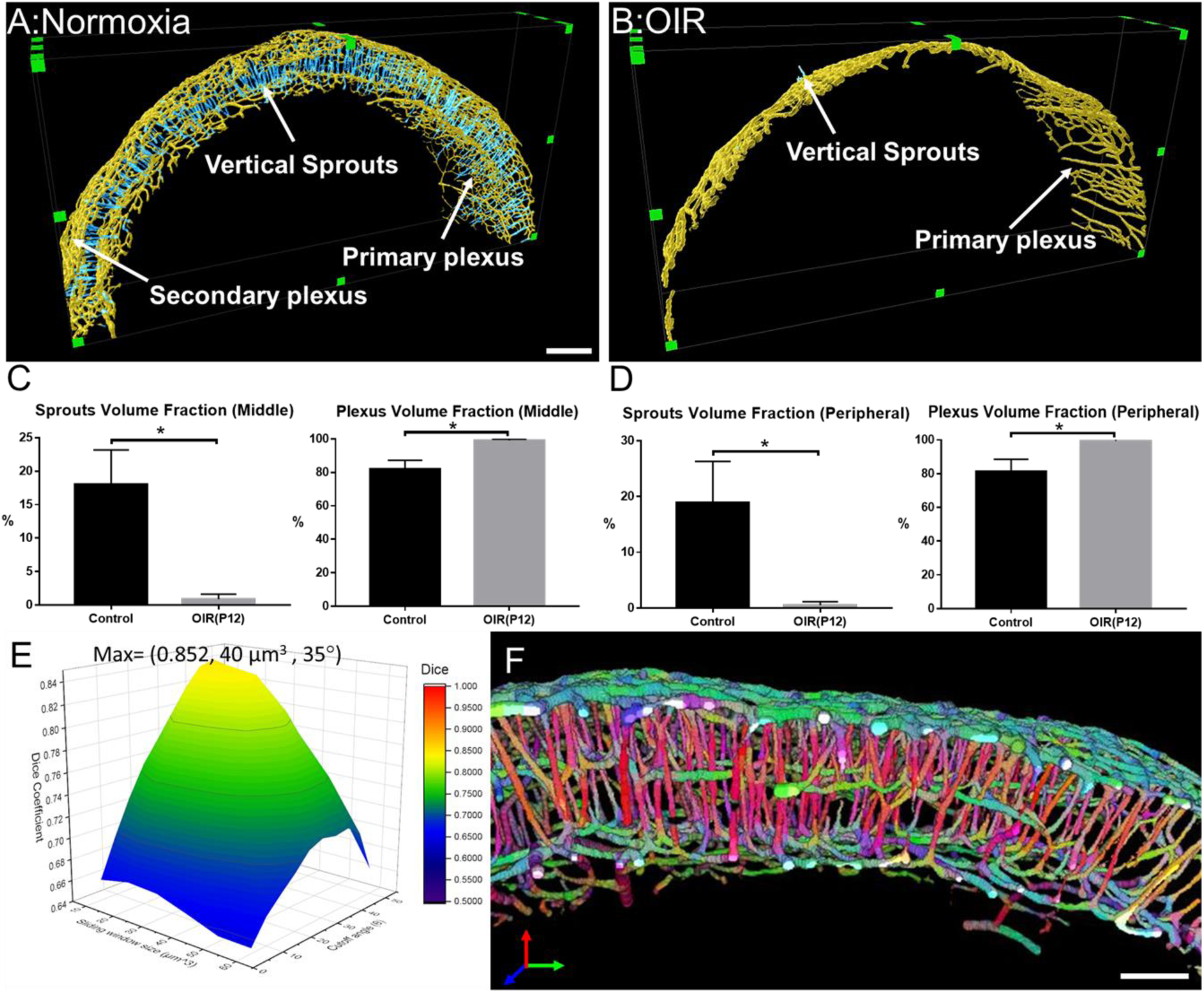
Quantitative analysis of the volumes for the vertical sprouts and plexuses. **(A-B)** Severe vaso-obliteration and depletion of vertical sprouts (in blue) and the secondary plexus (in yellow) were demonstrated in the 3-D rendering of the OIR group. **(C-D)** The quantification of the volume fraction of both sprouts and plexuses from both middle and peripheral regions indicate the significant vessel depletion in the OIR group (\**p* < 0.05 vs. Normoxia, n=5 per each group). **(E)** The surface plot of the dice coefficients on different sizes of sliding windows and values of cutoff angles. **(F)** The representative VOI demonstrating the orientation color-coded 3-D rendering of the vascular network (X: Blue, Y: Green: Z: Red). Scale bar: 200 µm for A-B and 100 µm for F.

## DISCUSSION

Over the past decades, advances in imaging modalities have dramatically enhanced the fundamental and experimental understanding of angiogenesis, treatment, and visual outcomes in numerous retinal disorder-mediated vasculopathies, including retinopathy of prematurity and diabetic retinopathy. In light of the paucity of findings in hyperoxia-induced micro-vascular injury, our light-sheet imaging uncovered the interconnection of the morphological network of vertical sprouts bridging the primary and secondary plexuses in a mouse model of oxygen-induced retinopathy (OIR). We developed a computational model to corroborate the reticular vascular network undergoing capillary obliteration, revealing preferential vaso-obliteration in the deep plexus and the bridging vessels under hyperoxia condition. Our laser light-sheet with dual illumination further revealed a 3-D drop-out phenomenon of the deep plexus preceding capillary obliteration in the superficial plexus, otherwise challenging with the 2-D imaging system^17,30^.

In the retina, vascular development is regulated by a host of molecular signaling pathways that are coordinated by interactions between external environmental cues and internal metabolic activities^2,3^. Murine retinal vasculature provides a viable platform to investigate epigenetic effects on angiogenesis^1,2,17^. Disturbances to the signaling pathways, including changes in oxygen and nutrient provision, promote pathological vaso-obliteration, resulting in structural and functional abnormalities during angiogenesis^1,6,17^. In diabetic retinopathy, hypoxia due to non-perfused capillary is a power inducer of vascular endothelial growth factor (VEGF) expression; thereby, promoting neo-angiogenesis and vascular permeability^7^. In retinopathy of prematurity, pathological depletion of capillary network progresses to irreversible blindness ^3,6^. To this end, developing LSFM imaging allows for imaging the 3-D hemispherical vascular network, uncovering multi-layer vascular changes, and quantifying epigenetic vs. genetic perturbations, with translational implication for therapeutic targets to restore vision ^2,17,31^.

The conventional modalities to image retinal vascular structures are usually prepared via flat-mount samples, with the fluorescently-labeled structures detected via confocal or fluorescence microscopy^18,19,32-35^. Unlike wide-field and confocal microscopes, LSFM enables rapid optical sectioning at high acquisition rates^22,23^. Optical sectioning generates a sheet of light, allowing for a selective plane to be illuminated and excited with high axial resolution; thereby, mitigating photobleaching^21^. Thus, LSFM imaging enables rapid imaging acquisition of the entire mouse retina in ∼2 minutes, minimizing the photo-toxicity while providing high throughput studies.

Advances in optical clearing techniques have rendered the rodent tissue or organ systems, including retinas, transparent for LSFM imaging^22,23,36-39^. These techniques remove lipids from tissues to minimize photon scattering while stabilizing the 3-D structural conformation ^40-42^, allowing for deep laser penetration into a large-sized samples^40,43-46^. 3-D imaging of murine hyperoxia-induced retinopathy often constitutes technical challenges in association with the premature ocular development and small size of the neonatal mouse eye. To elucidate the 3-D microvascular network in retinopathy of prematurity, we demonstrate the capacity of LSFM imaging to reveal the differential effects of pathologic hyperoxia in the multi-layers of the neonatal retinal vasculature. Our pipeline provides deep tissue penetration with high spatial resolution to visualize the reticular vascular network at different stages of retinal vasculopathies.

In human diabetic retinopathy, hypoxia-induced injury to the superficial and deep capillary plexus has been quantified via optical coherence tomography angiography (OCTA)^47,48^, branch retinal vein occusion^49^, and sickle cell disease^50^. For instance, OCTA shows vessel density from the deep capillary network as an early marker of diabetic vasculopathy^48^, otherwise challenging to detect with fundoscopic examination^47^. OCTA further demonstrates the reduction in vessel density within the superficial and deep capillary plexuses, that altered the corresponding retinal layers^51^. Although OCTA provides the clinical and in-vivo capability, the limitations of a relatively small field of view, and image artifacts due to patient movement/blinking underlie difficulties of capturing the entire retinal vasculature from the center to peripheral regions, especially in small animal models^52^. Thus, the aforementioned studies highlight the clinical relevance for elucidating the differential capillary depletion in a 3-D retinal vascular networks.

In the mouse model of oxygen-induced retinopathy, LSFM imaging, followed by computational analyses, quantifies the 3-D morphological and topological parameters for the entire vascular network. In addition to the vessel lengths, vessel volumes, and branching points, our computational analyses further quantified the global vs. local vascular connectivity by Euler numbers and clustering coefficients, respectively. Unlike the conventional estimation of the network connections via the branching points, the Euler numbers simultaneously reflect both connectivity and topological features in the 3-D network ^53-55^, facilitating the measurement of the connectivity of physiological structures, including the vascular networks and trabecular bones ^19,56-59^. In addition, the clustering coefficients capture the local connectivity and degree of organization of the regional vascular network for every vertex in the vascular and neuronal network^60-62^. Both Euler numbers and clustering coefficients enabled us to characterize the 3-D topography of the vascular network. Following hyperoxia-induced retinopathy, there was a significant reduction in global connectivity, but the local reticular pattern was preserved. We were also able to apply principal component analysis (PCA) to automatically segment the vertical sprouts from the plexuses and to quantify the volume fraction in the plexuses and vertical sprouts under normoxia vs. hyperoxia conditions, corroborating the differential depletion in vertical sprouts and the secondary plexus. Despite the well-defined formula to quantify the Euler numbers and clustering coefficients, the uncertainties reside in the accuracy of the segmentation of the blood vessels by using filament tracing. Machine learning-based automated segmentation for the vessels provides an alternative approach to abbreviate the segmentation process and to obviate the need for filament tracing. Taken together, we unraveled the preferential vaso-obliteration accompanied with the impaired vascular branching, connectivity, and reduced vessel volumes and lengths in OIR retinas.

Overall, LSFM illumination enables deep tissue penetration to unravel structural changes in 3-D retinal vascular networks. Integration of light-sheet imaging and 3-D computational analyses detects and quantifies global vaso-obliteration, but preserved local 3-D reticular vascular architecture in support of the differential vascular obliteration in hyperoxia-mediated retinopathy. With the copious data generated from advanced optics, quantitative strategy in 3-D, provides structural insights into retina disorders-associated vascular obliteration vs. proliferation, with translational significance for early detection and intervention to restore visual impairment.

## Material and Methods

### Ethics statement

All animal studies were performed in compliance with the IACUC protocol approved by the UCLA Office of Animal Research and in accordance with the guidelines set by the National Institutes of Health. Humane care and use of animals were observed to minimize distress and discomfort.

### The modified passive CLARITY for retina

We optimized the CLARITY methods for fixed retinas as previously described ^42,45^. The mice were euthanized at P12 for dissection of the ocular globe. We enucleated the intact ocular globe followed by fixation in 4% paraformaldehyde (PFA) (w/v) for 1.5 hours. We then removed the cornea, sclera, lens, and choroid while preserving the retina in its 3-D hemispherical configuration. The 3-D retinas were then immersed in 4% paraformaldehyde (w/v) overnight (Fig. 1D-➀), followed by overnight incubation in the monomer solution (4% Acrylamide (w/v), 0.05% Bis-Acrylamide (w/v), and 0.25% VA-044 initiator (w/v) in PBS) (Fig. 1D-➁). Next, samples were placed into a 37 °C water bath for 6 hours for hydrogel polymerization (Fig. 1D-➂). A degassing nitrogen flush or addition of an oil layer was performed to minimize exposure to oxygen. Next, retinas were incubated in the clear solution (4% w/v sodium dodecyl sulfate (SDS) and 1.25% w/v boric acid (pH 8.5) at 37 °C to remove non-transparent lipid contents. (Fig. 1D-➃). The retina was rinsed for an additional 24 hours in 1X PBS to remove residual SDS, followed by incubation in the refraction index matching solution RIMS (40 g histodenz in 30 ml of 0.02 M PBS with 0.1% tween-20 and 0.01% neutralized sodium azide (pH to 7.5 with NaOH)) to achieve transparency. This method enables preservation of the 3-D structure and maintains integrity of the retinal vasculature during clearing. Selected P12 retinas (n=3) were collected for direct immunostaining without optical clearing (non-cleared group) for comparison (Supplementary Fig.2).

### Animal model and OIR induction in postnatal mice

C57BL/6J mice were acquired from the UCLA Division of Laboratory and Animal Medicine colony. All mice were housed in 12:12 hour light-dark cycles. Female pregnant mice were fed *ad libitum* with standard rodent chow diet (Pico Lab Rodent Diet 20, cat#5053, Lab Diet, St. Louis, MO) and water during pregnancy and lactation. Oxygen-induced retinopathy (OIR) was produced using standard published guidelines^18^. Pups were designated at P0.5 on the morning that they were delivered. Per protocol recommendations, litters were culled to eight pups. Mothers were randomly assigned to normoxia or hyperoxia (OIR) conditions. To induce OIR, the newborn mice in OIR groups were exposed to 75% oxygen continuously in an airtight chamber (Biospherix Proox model 360, Parish, NY, USA) with their nursing mothers from P7 to P12 before removal to room air, whereas the normoxia group remained in room air (21% oxygen) throughout the postnatal period. Pups whose mothers demonstrated stress after removal from the oxygen chamber in the hyperoxia condition were fostered to a nursing mother until euthanasia at P12.

### Immunostaining for retinal vasculature

The optically transparent retinas were placed into a blocking buffer Perm/Block (1X PBS + 0.3% Triton-X + 0.2% bovine serum albumin + 5% of fetal bovine serum) solution for 1 hour at room temperature with gentle shaking. Primary biotinylated GS isolectin B4 (1:50, Vector lab, CA) was used to stain retinal vasculature. Following 2 days of incubation at 4 °C, retinas were washed with PBSTX (1X PBS + 0.3% Triton-X). Strepatavidin conjugated with Alexa-488 (1:100, Invitrogen, CA) was utilized to amplify primary-specific fluorescence. The retinas were washed with PBSTX (1X PBS + 0.3% Triton-X) following each step.

### Light-sheet fluorescence microscope (LSFM) with dual-illumination to image 3-D retinal vascular network

We custom-built our LSFM with dual-illumination for imaging the 3-D retinal vasculature ^21,22,63^. The detection arm is composed of a stereomicroscope (MVX10, Olympus, Japan) with a 1X magnification objective (NA = 0.25), a scientific CMOS (sCMOS, ORCA-Flash4.0 LT, Hamamatsu, Japan) and a series of filters (Exciter: FF01-390/482/532/640; Emitter: FF01-446/510/581/703; Dichroic: Di01-R405/488/532/635, Semrock, New York, USA). The illumination arm, orthogonal to detection, is composed of the continuous wave laser at 473 nm and 532 nm (LMM-GBV1-PF3-00300-05, Laserglow Technologies, Canada) (Supplementary Figures 3A-B). The lateral and axial resolutions are 5 µm and 6 µm, respectively (Supplementary Figure 3C). For image acquisition, the samples were immersed in the RIMS (RI: 1.46–1.48) with a 1% agarose solution in a Borosilicate glass tube (RI = 1.47, Pyrex 7740, Corning, New York, USA) to reduce refraction and reflection from various interfaces. The glass sample holder was placed in a 3-D-printed opening chamber, made of acrylonitrile butadiene styrene (ABS) (uPrint, Stratasys, USA) and was filled with 99.5% glycerol (RI = 1.47).

### Computational analyses for the morphological and topological parameters of the vascular network

The raw data were preprocessed in ImageJ to remove stationary noise^64,65^ and the background was reduced by rolling ball background subtraction (20 pixels in radius). To provide the axial visualization or the depth of the 3-D vascular network, we applied the depth-coded plugin in ImageJ to enhance visualization of the superficial and deep capillaries. In addition, 3-D rendering and semi-automated filament tracing were performed and processed in Amira 6.1. The results of filament tracing were used to quantify both the morphological (Euler-Poincaré characteristic) and topological parameters (clustering coefficients). The Euler number, χ, of the 3-D object is defined as follows^53,55^:

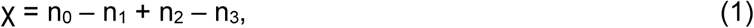

where n_0,_ n_1,_ n_2,_ and n_3_ are the numbers of vertices (V), edges (E), faces (F) and the individual voxels contained in a 3-D object, respectively. In the setting of increasing connected edges, the Euler number decreases while the network becomes well-connected. We provided an analysis of the changes in Euler number in response to the morphological difference in Supplementary Figure 4A. The Euler number remained the same when the new connections form a branching structure (first row of Supplementary Figure 4A). In addition, the Euler number was reduced when a new connection formed loops in a reticular-like structure (second row in Supplementary Figure 4A), whereas the Euler number was increased when the new connection formed the disconnected objects (third row in Supplementary Figure 4A). Thus, two factors are the main contributors to the Euler number: 1) the number of loops (holes), and 2) the numbers of disconnected objects^17^. In this context, we defined the connectivity for the vascular network as ^56^:

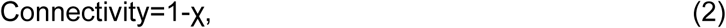

In comparison to the Euler number, the clustering coefficient (C) reflects the local connectivity of individual vertex or node in the vasculature ^60-62,66^. The clustering coefficient of each vertex and the average clustering coefficient C are defined as follows, respectively:

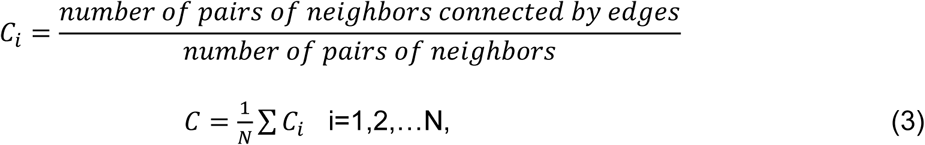

The local clustering coefficient *C*_i_ for a vertex v_i_ is given by the proportion of links between the vertices within its neighborhood divided by all possible connections between its neighbors (Supplementary Figure 4B). For detailed mathematical calculation, a graph G (Equation 4) consists of a set of edges, ***E***, and vertices, ***V***, and an edge, e_ij_, is connecting vertex v_i_ with vertex v_j_ (Supplementary Figure 4B). We define the neighborhood set *N*_i_ of vertex v_i_ for its directly connected neighbors (Equation 5). By defining k_i_=| ***N***_i_ | as the number of neighbors in set ***N***_i_ for vertex v_i_, the clustering coefficient C_i_ for vertex v_i_ can be calculated (Equation 6).

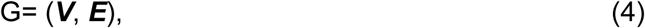

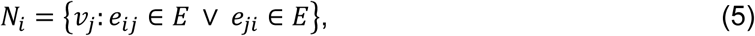

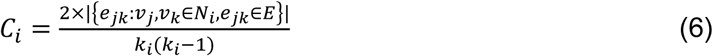

### Quantitative Analysis of vascular plexuses and vertical sprouts in retinal vasculature

The segmented 3D image stacks were derived from the filament tracing results for intact retinal vasculature, and they were resampled along the axis passing through and being perpendicular to the optical disk. To quantify the volume of vertical sprouts, we developed automated segmentation based on principal component analysis (PCA) for reducing the dimensionality for the large data set. Considering a thin section from the retinal vasculature image stack with the origin at the center of mass, the vessels belonging to the primary and secondary plexuses are mainly in azimuthal or polar directions, whereas the majority of the vertical sprouts develop in a radial direction (Supplementary Figure 1A). To quantify the vessel direction for 3-D image stacks, we introduced a sliding window to sample the principal component representing the vessel orientation inside the window (Supplementary Figure 1A). The coordinates of each segmented blood vessel inside the sliding window were extracted and recorded for deriving the representative principal component for each window (Supplementary Figure 1A). Next, we applied the center of the mass as the origin to calculate the angle (θ) between the principal vector of the voxel and the vector from the origin to the center of the voxel (Supplementary Figure 1A). We have optimized the sliding window size and the value of the cut-off angle for differentiating the vertical sprouts from the vascular plexuses.

### Statistical analysis

All data were presented as mean ± SD. Statistical significance was using unpaired two-tailed Student’s t-test for comparison of two groups and one-way ANOVA with Tukey post hoc analysis for multiple group comparisons. The level of significance was set at p < 0.05.

## Acknowledgment

The present work was funded by NIH grants National Institutes of Health R01HL083015 (TKH), R01HL111437 (TKH), R01HL129727 (TKH), R01HL118650 (TKH), VA MERIT AWARD I01 BX004356 (TKH), K99 HL148493 (YD) and AHA 18CDA34110338 (YD).

## Author contributions

C.C, A.C, and T.K.H conceived the research and design the experiment with the input from J.J.Z, K.I.B, Y.D, L.K.G, and S.L. C.C, S.M, P.A and V.G developed the quantification method and performed the analysis. A.C, L.K.G, and S.L provided the OIR and normoxia mouse model. M.M.S, P.A, K.B, X.D, and P.G dissected and processed the retina samples for imaging. C.C performed the imaging experiment and build up the imaging system with Y.D. C.C, A.C, and T.K.H wrote the manuscript, with contributions from all authors. All authors reviewed the manuscript.

## Data availability

The authors declare that the main data supporting the findings of this study are available within the article and its Supporting Information files. Extra data are available from the corresponding author on a reasonable request.

## Code availability

Code that supports the findings of this study is available upon reasonable request from the first author but third-party restrictions may apply (e.g., MATLAB, Amira).

## Supplementary Notes

**Supplementary movie 1. The 3-D rendering of the vascular network to demonstrate the vertical sprouts embedded between primary and secondary plexus.**

**Supplementary movie 2. The filament tracing result of the intact control retinal vascular network.**

**Supplementary movie 3. The 3-D rendering of the intact control retinal vascular network.**

**Supplementary movie 4. The 3-D rendering of the intact OIR retinal vascular network.**

**Supplementary movie 5. The result of filament tracing from the intact OIR retinal vascular network.**

### Representative results of integration of LSFM and tissue-clearing

The representative images for the maximum intensity projection from LSFM were compared between the passive CLARITY-treated retina and control, demonstrating the capacity of optical clearing to uncover the microvasculature that would otherwise be challenging with conventional imaging modalities. Note the pre and post-optical clarity ability to image the deep vascular plexus (Supplementary Figure 2B).

### Principal component analysis (PCA) of vasculature

PCA is a mathematical process to reduce the dimensions and unravel the important characteristics of the data. It is a linear transformation calculating the new set of the coordinate system for the data such that the greatest variation was introduced to the projected data points to the new axis1. The ***r***i,j (i=1,2…N, j=1-3) represents the coordinate vectors from the vasculature (Equation 1). N here represents the numbers of vessel elements inside the sliding window while j represents 3-D space dimensions (X, Y, and Z). The new coordinates are computed from the covariance matrix (Equation 1,2) of the data and applied to the eigenvector decomposition (Equation 3,4) process to derive the new axes called principal components (PC) i.e. The PC1 is the eigenvector with the largest possible variance and each succeeding component such as PC2 and PC3 has the highest variance possible under the constraint that it is orthogonal to the preceding components. Lastly, the angle between vectors is calculated through the relation of the inner product (Equation 5).

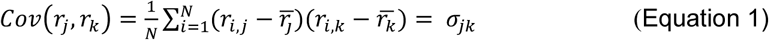

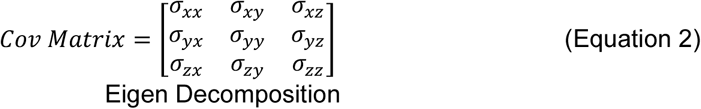

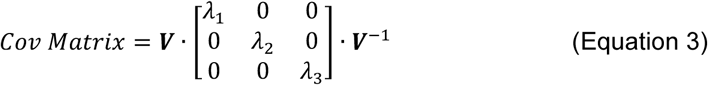

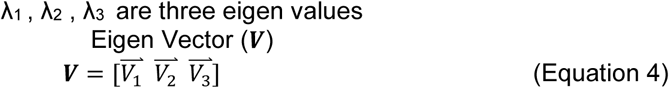

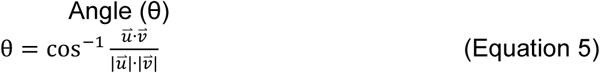

---

**Supplementary Figure 1.**
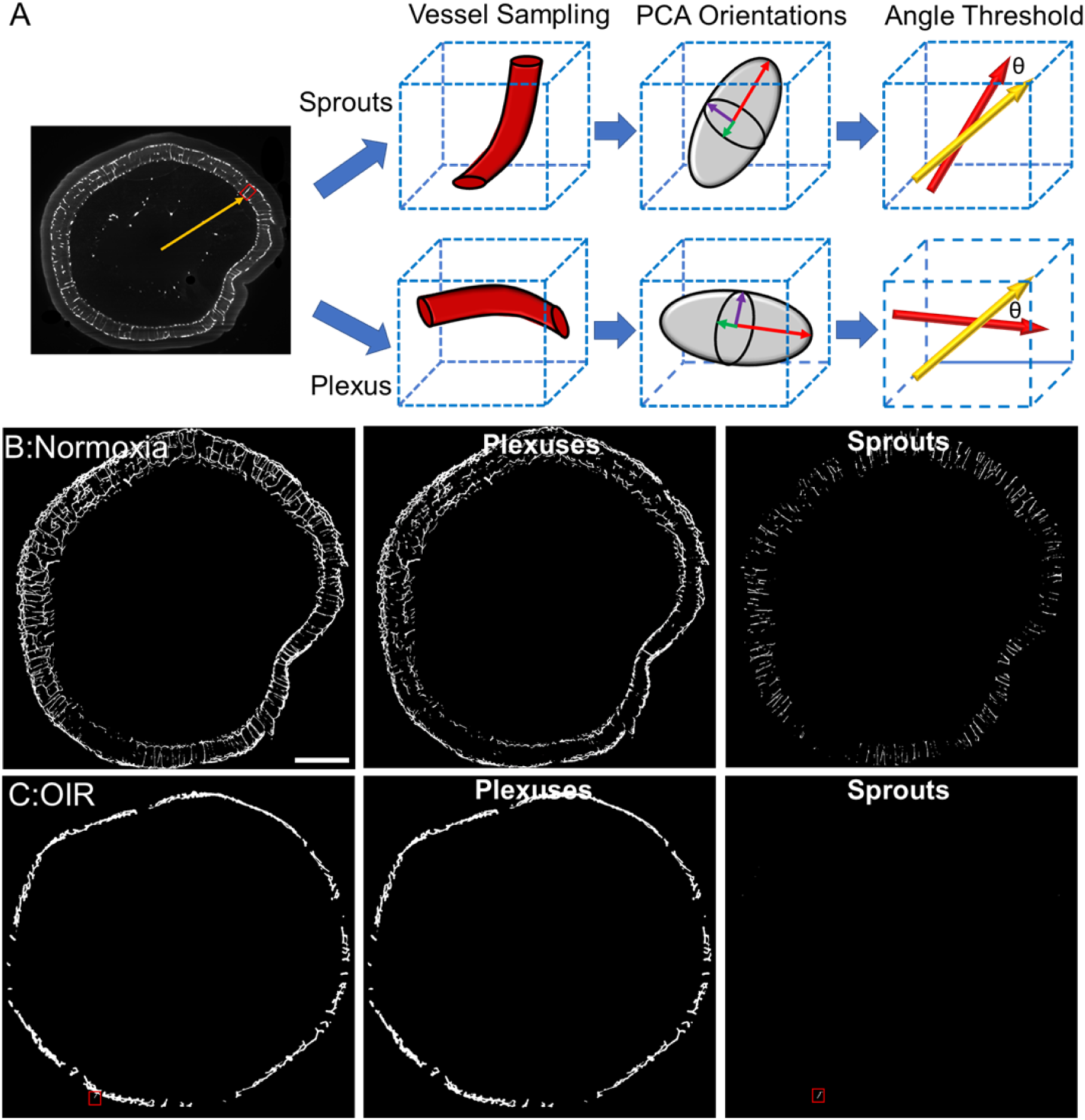
Schematic illustration and result of the automated segmentation for the vertical sprouts and plexuses. **(A)** The schematic plot indicates the distinct characteristics between the vertical sprouts and plexus serving as the bases for the automated segmentation method. **(B-C)** The representative MIP images from the image stacks demonstrated the separation between the vertical sprouts and plexus from automated segmentation for both normoxia and OIR (sprout highlighted in a red box) groups. Scale bar: 500 µm.

**Supplementary Figure 2.**
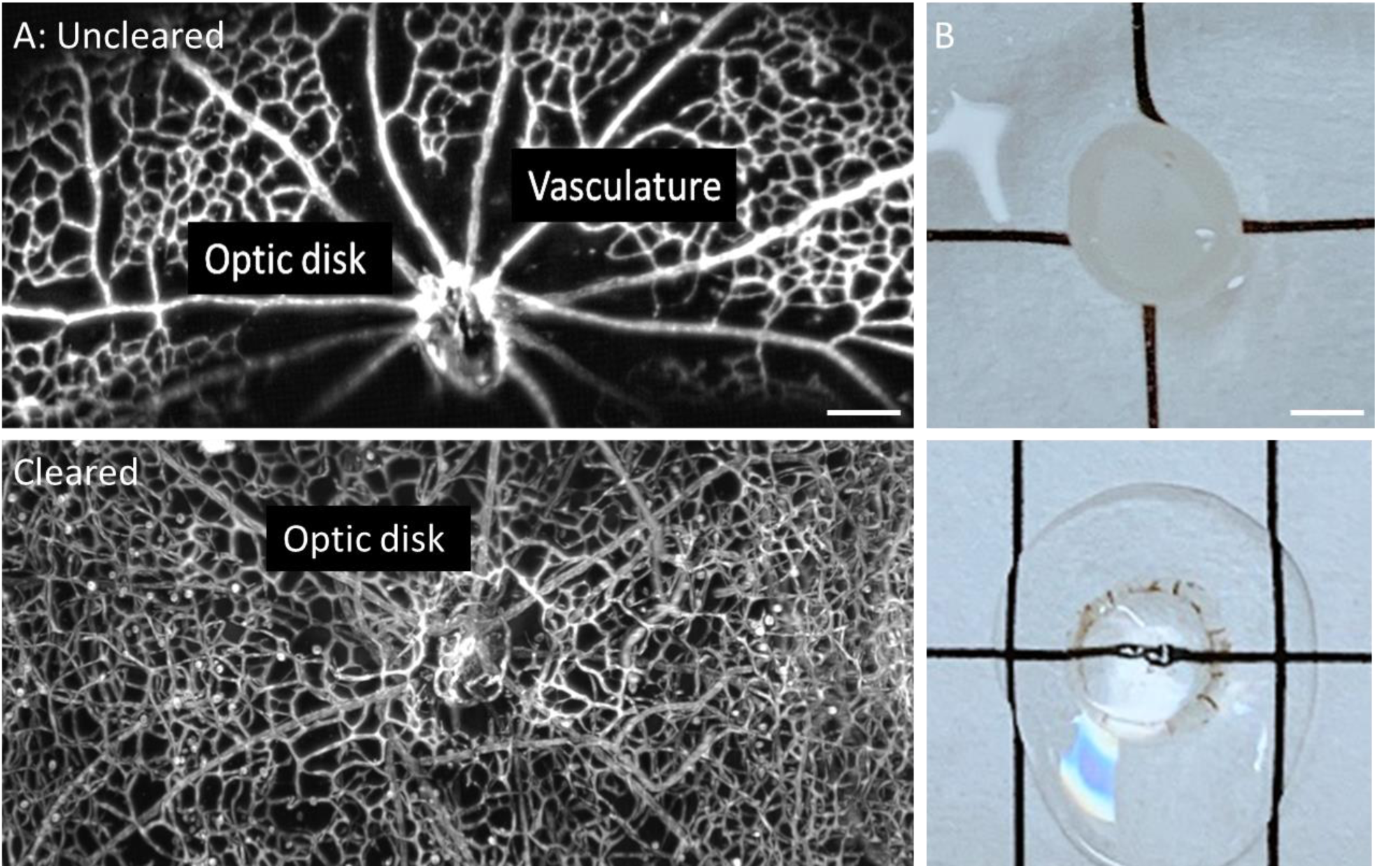
Representative comparison of retinas with and without tissue clearing. **(A)** The detailed vasculature post CLARITY method was demonstrated by maximum intensity projection in comparison to the uncleared retina. **(B)** Mouse retinas at postnatal 12 days prior to (top) and post (bottom) modified CLARITY treatment. Scale bar: (A) 100 µm; (B) 1 mm

**Supplementary Figure 3.**
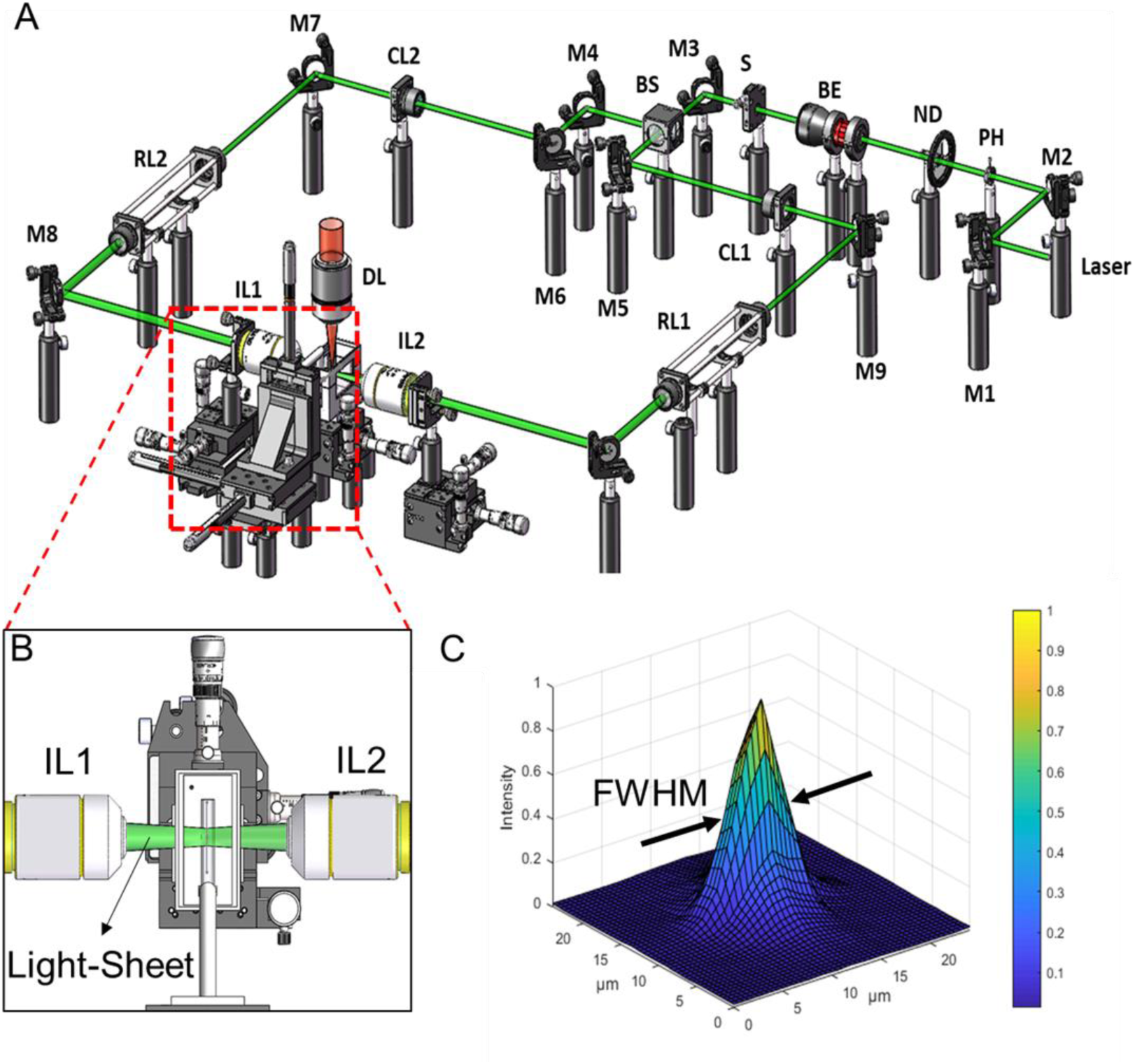
The schematic diagram of the dual-illumination LSFM. **(A)** The 3-D schematic highlights the individual optical components of the LSFM system. M1-10: mirror; PH: pinhole; ND: neutral density filter; BE: beam expander; S: slit; CL1-2: cylindrical lens; RL1-2: Relay lens; IL1-2: illumination lens; DL: detection lens. **(B)** Dual-sided illumination covers the retina in the chamber. **(C)** Point spread function (PSF) of the 0.53 µm fluorescent bead in the lateral direction.

**Supplementary Figure 4.**
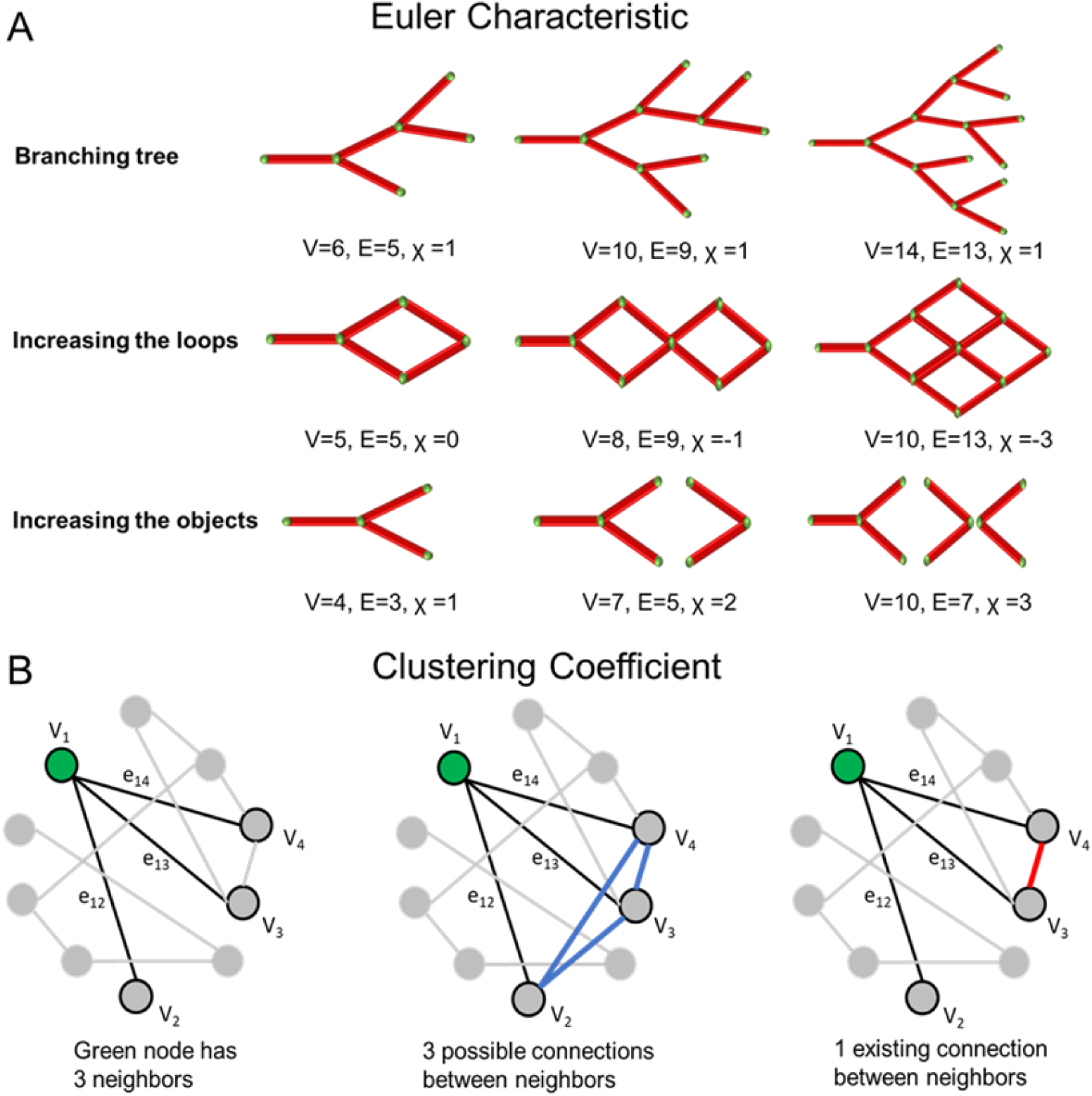
Schematic representation to quantify the Euler characteristic and clustering coefficients. **(A)** Euler characteristic χ for the vascular network is determined by the numbers of loops and objects rather than branching points. Both factors generate new nodes and edges unequally, and both increase the difference in the formula. The entire vascular network is leaning toward the reticular-like in structure, resulting in a reduction in χ value. V: Vertices, E: edges. **(B)** The definition of the clustering coefficient for a specific node (green) is calculated by dividing the existing links (red) over all possible links (blue) among neighbors, that is, the representative value of the schematic is 0.33

**Table 1.**
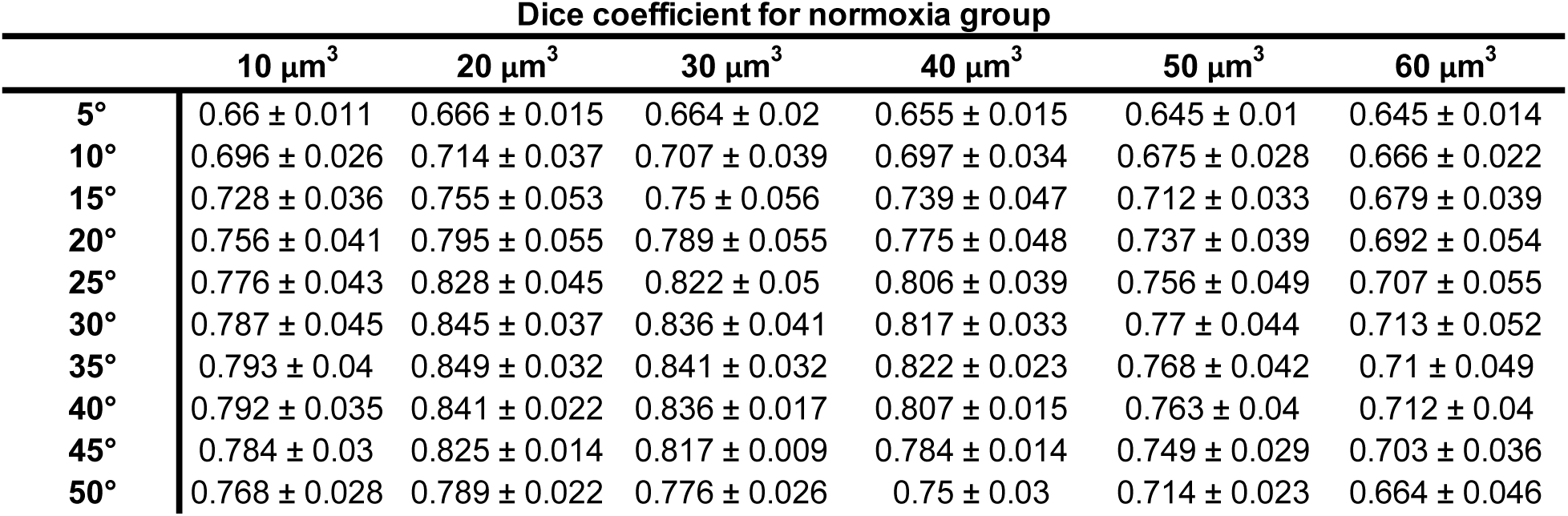
Average dice coefficients of various sliding window sizes and cutoff angles of the control group.

**Table 2.**
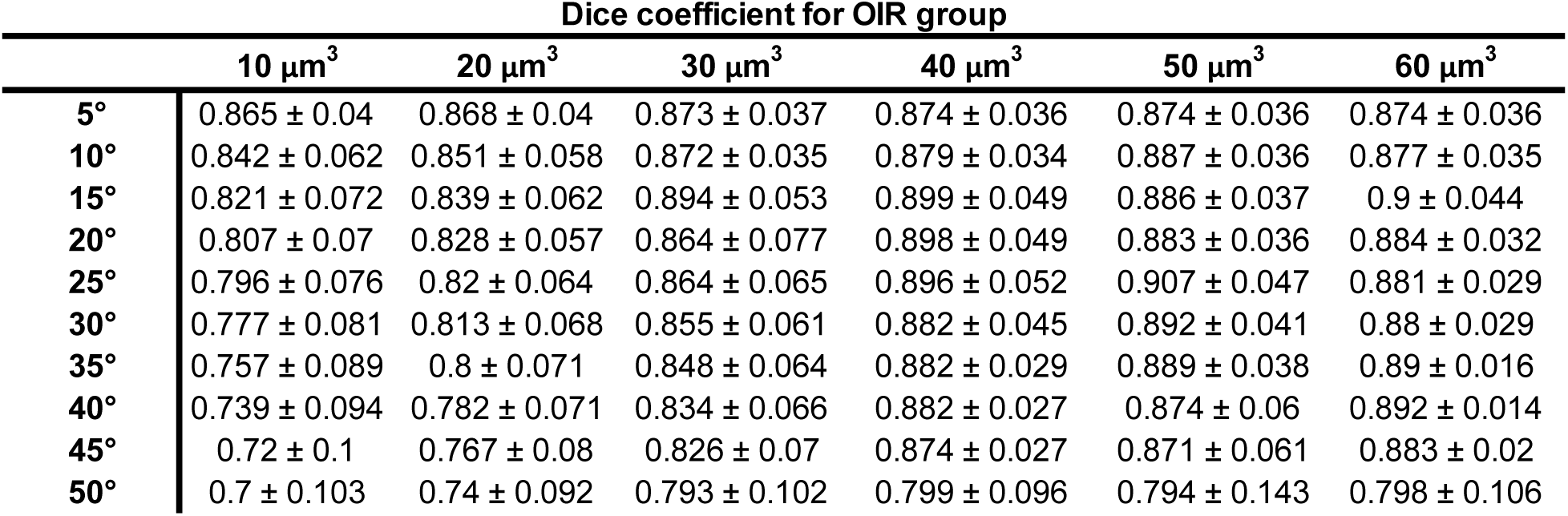
Average dice coefficients of various sliding window sizes and cutoff angles of the OIR group.

**Table 3.** Average dice coefficients of various sliding window sizes and cutoff angles of both groups.

| Dice coefficient for both groups |  |  |  |  |  |  |
| --- | --- | --- | --- | --- | --- | --- |
| | 10 $\mu\text{m}^3$ | 20 $\mu\text{m}^3$ | 30 $\mu\text{m}^3$ | 40 $\mu\text{m}^3$ | 50 $\mu\text{m}^3$ | 60 $\mu\text{m}^3$ |
| 5° | 0.763 ± 0.113 | 0.767 ± 0.111 | 0.769 ± 0.115 | 0.765 ± 0.12 | 0.76 ± 0.125 | 0.76 ± 0.125 |
| 10° | 0.769 ± 0.088 | 0.783 ± 0.085 | 0.789 ± 0.095 | 0.788 ± 0.102 | 0.781 ± 0.117 | 0.771 ± 0.116 |
| 15° | 0.775 ± 0.069 | 0.797 ± 0.068 | 0.822 ± 0.092 | 0.819 ± 0.096 | 0.799 ± 0.098 | 0.79 ± 0.124 |
| 20° | 0.782 ± 0.057 | 0.811 ± 0.054 | 0.826 ± 0.071 | 0.836 ± 0.079 | 0.81 ± 0.085 | 0.788 ± 0.111 |
| 25° | 0.786 ± 0.054 | 0.824 ± 0.049 | 0.843 ± 0.056 | 0.851 ± 0.063 | 0.831 ± 0.092 | 0.794 ± 0.102 |
| 30° | 0.782 ± 0.057 | 0.829 ± 0.05 | 0.846 ± 0.047 | 0.849 ± 0.049 | 0.831 ± 0.076 | 0.796 ± 0.098 |
| 35° | 0.775 ± 0.062 | 0.824 ± 0.054 | 0.845 ± 0.044 | 0.852 ± 0.04 | 0.829 ± 0.074 | 0.8 ± 0.103 |
| 40° | 0.765 ± 0.066 | 0.812 ± 0.054 | 0.835 ± 0.04 | 0.844 ± 0.044 | 0.819 ± 0.074 | 0.802 ± 0.1 |
| 45° | 0.752 ± 0.07 | 0.796 ± 0.056 | 0.821 ± 0.041 | 0.829 ± 0.052 | 0.81 ± 0.077 | 0.793 ± 0.101 |
| 50° | 0.734 ± 0.073 | 0.764 ± 0.061 | 0.784 ± 0.063 | 0.775 ± 0.065 | 0.754 ± 0.094 | 0.731 ± 0.1 |

## References

1 Liu, C.-H., Wang, Z., Sun, Y. & Chen, J. Animal models of ocular angiogenesis: from development to pathologies. The FASEB Journal 31, 4665–4681 (2017).

2 Selvam, S., Kumar, T. & Fruttiger, M. Retinal vasculature development in health and disease. Progress in retinal and eye research 63, 1–19 (2018).

3 Fruttiger, M. Development of the retinal vasculature. Angiogenesis 10, 77–88 (2007).

4 Zin, A. & Gole, G. A. Retinopathy of prematurity-incidence today. Clinics in perinatology 40, 185–200 (2013).

5 Cavallaro, G. et al. The pathophysiology of retinopathy of prematurity: an update of previous and recent knowledge. Acta ophthalmologica 92, 2–20 (2014).

6 Gariano, R. F. & Gardner, T. W. Retinal angiogenesis in development and disease. Nature 438, 960 (2005).

7 Semeraro, F., Cancarini, A., Rezzola, S., Romano, M. & Costagliola, C. Diabetic retinopathy: vascular and inflammatory disease. Journal of diabetes research 2015 (2015).

8 Antonetti, D. A. et al. Diabetic retinopathy: seeing beyond glucose-induced microvascular disease. Diabetes 55, 2401–2411 (2006).

9 Lee, D.-H. et al. Risk factors for retinal microvascular impairment in type 2 diabetic patients without diabetic retinopathy. PloS one 13, e0202103 (2018).

10 Chui, T. Y. et al. In Frontiers in Optics. FM3F. 1 (Optical Society of America).

11 Shah, P. K. et al. Retinopathy of prematurity: past, present and future. World journal of clinical pediatrics 5, 35 (2016).

12 Talisa, E. et al. Detection of microvascular changes in eyes of patients with diabetes but not clinical diabetic retinopathy using optical coherence tomography angiography. Retina 35, 2364–2370 (2015).

13 Dimitrova, G., Chihara, E., Takahashi, H., Amano, H. & Okazaki, K. Quantitative retinal optical coherence tomography angiography in patients with diabetes without diabetic retinopathy. Investigative ophthalmology & visual science 58, 190–196 (2017).

14 D’Amato, G. et al. Sequential Notch activation regulates ventricular chamber development. Nature cell biology 18, 7–20, doi:10.1038/ncb3280 (2016).

15 Takase, N. et al. Enlargement of foveal avascular zone in diabetic eyes evaluated by en face optical coherence tomography angiography. Retina 35, 2377–2383 (2015).

16 Cao, D. et al. Optical coherence tomography angiography discerns preclinical diabetic retinopathy in eyes of patients with type 2 diabetes without clinical diabetic retinopathy. Acta diabetologica 55, 469–477 (2018).

17 Stahl, A. et al. The mouse retina as an angiogenesis model. Investigative ophthalmology & visual science 51, 2813–2826 (2010).

18 Connor, K. M. et al. Quantification of oxygen-induced retinopathy in the mouse: a model of vessel loss, vessel regrowth and pathological angiogenesis. Nature protocols 4, 1565 (2009).

19 Milde, F., Lauw, S., Koumoutsakos, P. & Iruela-Arispe, M. L. The mouse retina in 3D: quantification of vascular growth and remodeling. Integrative Biology 5, 1426–1438 (2013).

20 Henning, Y., Osadnik, C. & Malkemper, E. P. EyeCi: Optical clearing and imaging of immunolabeled mouse eyes using light-sheet fluorescence microscopy. Experimental eye research 180, 137–145 (2019).

21 Ding, Y. et al. Multiscale light-sheet for rapid imaging of cardiopulmonary system. JCI insight 3 (2018).

22 Baek, K. I. et al. Advanced microscopy to elucidate cardiovascular injury and regeneration: 4D light-sheet imaging. Progress in biophysics and molecular biology 138, 105–115 (2018).

23 Power, R. M. & Huisken, J. A guide to light-sheet fluorescence microscopy for multiscale imaging. Nature methods 14, 360 (2017).

24 McDole, K. et al. In toto imaging and reconstruction of post-implantation mouse development at the singlecell level. Cell 175, 859-876. e833 (2018).

25 Ding, Y. et al. Integrating light-sheet imaging with virtual reality to recapitulate developmental cardiac mechanics. JCI insight 2 (2017).

26 Pawley, J. Handbook of biological confocal microscopy. (Springer Science & Business Media, 2010).

27 Stelzer, E. H. Light-sheet fluorescence microscopy for quantitative biology. Nat. Methods 12, 23–26 (2015).

28 Weinhaus, R. S., Burke, J. M., Delori, F. C. & Snodderly, D. M. Comparison of fluorescein angiography with microvascular anatomy of macaque retinas. Experimental eye research 61, 1–16 (1995).

29 Sudre, C. H., Li, W., Vercauteren, T., Ourselin, S. & Cardoso, M. J. In Deep learning in medical image analysis and multimodal learning for clinical decision support 240–248 (Springer, 2017).

30 Calzi, S. L. et al. Progenitor cell combination normalizes retinal vascular development in the oxygen-induced retinopathy (OIR) model. JCI insight 4 (2019).

31 Chang, B. In Retinal Degeneration 27–39 (Springer, 2012).

32 Kim, J. et al. YAP/TAZ regulates sprouting angiogenesis and vascular barrier maturation. The Journal of clinical investigation 127, 3441–3461 (2017).

33 Yoon, C.-H. et al. Diabetes-induced Jagged1 overexpression in endothelial cells causes retinal capillary regression in a murine model of diabetes mellitus: Insights into diabetic retinopathy. Circulation 134, 233–247 (2016).

34 Wilhelm, K. et al. FOXO1 couples metabolic activity and growth state in the vascular endothelium. Nature 529, 216 (2016).

35 Dubrac, A. et al. Nck-dependent pericyte migration promotes pathological neovascularization in ischemic retinopathy. Nature communications 9 (2018).

36 Ding, Y. et al. In Advanced Biomedical and Clinical Diagnostic and Surgical Guidance Systems XVI. 104841C (International Society for Optics and Photonics).

37 Lee, J. et al. 4-Dimensional light-sheet microscopy to elucidate shear stress modulation of cardiac trabeculation. The Journal of clinical investigation 126, 1679–1690 (2016).

38 Prahst, C. et al. Mouse retinal cell behaviour in space and time using light sheet fluorescence microscopy. Elife 9, e49779 (2020).

39 Singh, J. N., Nowlin, T. M., Seedorf, G. J., Abman, S. H. & Shepherd, D. P. Quantifying three-dimensional rodent retina vascular development using optical tissue clearing and light-sheet microscopy. Journal of biomedical optics 22, 076011 (2017).

40 Richardson, D. S. & Lichtman, J. W. Clarifying tissue clearing. Cell 162, 246–257 (2015).

41 Tomer, R., Ye, L., Hsueh, B. & Deisseroth, K. Advanced CLARITY for rapid and high-resolution imaging of intact tissues. Nature protocols 9, 1682 (2014).

42 Sung, K. et al. Simplified three-dimensional tissue clearing and incorporation of colorimetric phenotyping. Scientific reports 6, 30736 (2016).

43 Gradinaru, V., Treweek, J., Overton, K. & Deisseroth, K. Hydrogel-tissue chemistry: Principles and applications. Annual review of biophysics 47, 355–376 (2018).

44 Richardson, D. S. & Lichtman, J. W. SnapShot: tissue clearing. Cell 171, 496-496. e491 (2017).

45 Chung, K. et al. Structural and molecular interrogation of intact biological systems. Nature 497, 332 (2013).

46 Renier, N. et al. iDISCO: a simple, rapid method to immunolabel large tissue samples for volume imaging. Cell 159, 896–910 (2014).

47 Sousa, D. C. et al. Optical coherence tomography angiography study of the retinal vascular plexuses in type 1 diabetes without retinopathy. Eye, 1-5 (2019).

48 Zhu, T. P. et al. COMPARISON OF PROJECTION-RESOLVED OPTICAL COHERENCE TOMOGRAPHY ANGIOGRAPHY-BASED METRICS FOR THE EARLY DETECTION OF RETINAL MICROVASCULAR IMPAIRMENTS IN DIABETES MELLITUS. Retina (Philadelphia, Pa.) (2019).

49 Kim, J. T., Chun, Y. S., Lee, J. K., Moon, N. J. & Yi, D. Y. Comparison of Vessel Density Reduction in the Deep and Superficial Capillary Plexuses in Branch Retinal Vein Occlusion. Ophthalmologica, 1–9 (2019).

50 Ong, S. S. et al. Retinal Thickness and Microvascular Changes in Children With Sickle Cell Disease Evaluated by Optical Coherence Tomography (OCT) and OCT Angiography. American journal of ophthalmology (2019).

51 Lavia, C. et al. Reduced vessel density in the superficial and deep plexuses in diabetic retinopathy is associated with structural changes in corresponding retinal layers. PloS one 14 (2019).

52 De Carlo, T. E., Romano, A., Waheed, N. K. & Duker, J. S. A review of optical coherence tomography angiography (OCTA). International journal of retina and vitreous 1, 5 (2015).

53 Legland, D., Kiêu, K. & Devaux, M.-F. Computation of Minkowski measures on 2D and 3D binary images. Image Analysis & Stereology 26, 83–92 (2011).

54 Nagel, W., Ohser, J. & Pischang, K. An integral-geometric approach for the Euler-Poincaré characteristic of spatial images. Journal of microscopy 198, 54–62 (2000).

55 Toriwaki, J. & Yonekura, T. Euler number and connectivity indexes of a three dimensional digital picture. FORMA-TOKYO- 17, 183–209 (2002).

56 Nyengaard, J. R. Stereologic methods and their application in kidney research. Journal of the American Society of Nephrology 10, 1100–1123 (1999).

57 Willführ, A. et al. Estimation of the number of alveolar capillaries by the Euler number (Euler-Poincaré characteristic). American Journal of Physiology-Lung Cellular and Molecular Physiology 309, L1286–L1293 (2015).

58 Muehlfeld, C. Quantitative morphology of the vascularisation of organs: a stereological approach illustrated using the cardiac circulation. Annals of Anatomy-Anatomischer Anzeiger 196, 12–19 (2014).

59 Odgaard, A. & Gundersen, H. Quantification of connectivity in cancellous bone, with special emphasis on 3-D reconstructions. Bone 14, 173–182 (1993).

60 Amat-Roldan, I., Berzigotti, A., Gilabert, R. & Bosch, J. Assessment of hepatic vascular network connectivity with automated graph analysis of dynamic contrast-enhanced US to evaluate portal hypertension in patients with cirrhosis: a pilot study. Radiology 277, 268–276 (2015).

61 Czech, W., Dzwinel, W., Goryczka, S., Arodz, T. & Dudek, A. Z. Exploring complex networks with graph investigator research application. Computing and Informatics 30, 381–410 (2012).

62 Bullmore, E. & Sporns, O. Complex brain networks: graph theoretical analysis of structural and functional systems. Nature reviews neuroscience 10, 186 (2009).

63 Ding, Y. et al. Light-sheet fluorescence imaging to localize cardiac lineage and protein distribution. Scientific reports 7, 42209 (2017).

64 Fehrenbach, J., Weiss, P. & Lorenzo, C. Variational algorithms to remove stationary noise: applications to microscopy imaging. IEEE transactions on image processing 21, 4420–4430 (2012).

65 Fehrenbach, J. & Weiss, P. Processing stationary noise: Model and parameter selection in variational methods. SIAM Journal on Imaging Sciences 7, 613–640 (2014).

66 Watts, D. J. & Strogatz, S. H. Collective dynamics of ‘small-world’networks. nature 393, 440 (1998).

## References

1 Jolliffe, I. T. & Cadima, J. Principal component analysis: a review and recent developments. Philosophical Transactions of the Royal Society A: Mathematical, Physical and Engineering Sciences 374, 20150202 (2016).

